# Broad frequency sensitivity and complex neural coding in the larval zebrafish auditory system

**DOI:** 10.1101/2020.09.17.301242

**Authors:** Rebecca E. Poulsen, Leandro A. Sholz, Lena Constantin, Itia Favre-Bulle, Gilles C. Vanwalleghem, Ethan K. Scott

## Abstract

Most animals have complex auditory systems that identify salient features of the acoustic landscape to direct appropriate responses. In fish, these features include the volume, frequency, complexity, and temporal structure of auditory stimuli transmitted through water. Larval fish have simple brains compared to adults, but swim freely and depend on sophisticated sensory processing for survival. Zebrafish larvae, an important model for studying brain-wide neural networks, have thus far been found to possess a rudimentary auditory system, sensitive to a narrow range of frequencies and without evident sensitivity to auditory features that are salient and ethologically important to adult fish. Here, we have combined a novel method for delivering water-borne sounds, a diverse assembly of acoustic stimuli, and whole-brain calcium imaging to describe the responses of individual auditory neurons across the brains of zebrafish larvae. Our results reveal responses to frequencies ranging from 100Hz to 4kHz, with evidence of frequency discrimination from 100Hz to 2.5kHz. Frequency-selective neurons are located in numerous regions of the brain, and neurons responsive to the same frequency are spatially grouped in some regions. Using functional clustering, we identified categories of neurons that are selective for pure tones of a single frequency, white noise, the sharp onset of auditory stimuli, and stimuli involving a gradual crescendo. These results suggest a more nuanced auditory system than has previously been described in larval fish and provide insights into how a young animal’s auditory system can both function acutely and serve as the scaffold of a more complex adult system.

## RESULTS AND DISCUSSION

The auditory systems of fish are crucial for their survival, informing behaviors that include escaping from predators, searching for food, and communicating with each other[1–5]. There are several properties of sound stimuli that allow fish to extract ethologically important information, and therefore, to mount appropriate behavioral responses. These include the sounds’ frequencies, amplitudes, durations, and whether they are pure tones of a single frequency or complex sounds composed of multiple frequencies. Various fish species have been shown to respond behaviorally to a range of different auditory stimulus properties [2, 6–9], and some of the sensory neurons mediating these behaviors have been characterized electrophysiologically [10–14]. Nonetheless, our understanding of these auditory networks remains incomplete in terms of the categories of responsive neurons, their locations and connections, and the ways in which they work together to process auditory information.

Zebrafish larvae, which provide unparalleled opportunities for the description of sensory circuits and networks [15–18], have thus far been reported to have a rudimentary auditory system [19, 20]. As a result, many of the circuit-level details of complex auditory processing in fish remain unexplored, and it remains unclear whether the larvae of any fish species extract complex properties from auditory stimuli to inform behavior [21–23]. In this study, we performed an analysis of larval zebrafish hearing using whole-brain calcium imaging at cellular resolution while applying controlled auditory stimuli with a range of amplitudes, frequencies, complexities, and durations. Because our approach was both comprehensive (spanning the entire brain) and detailed (resolving individual neurons), it offered the potential to reveal brain-wide auditory networks in a way that has not previously been possible, as well as shed light on the auditory capabilities of an aquatic species has during the early stages of development.

Our approach required the accurate and reproducible delivery of sound [24], but the air-water interface complicates this stimulation in aquatic systems, especially in cases where underwater speakers are impractical. Past studies used different approaches for delivering sound to zebrafish larvae [7, 25–27], and these larvae have variously been shown to respond behaviorally to frequencies up to 200Hz [11, 26], 400hz [28], or 1000-1200Hz [7, 25, 29]. The first calcium imaging study of larval zebrafish hearing, using an air speaker, found strong neural responses in the 100-400Hz range, with weak responses up to 800Hz [19]. A more recent imaging study, using an array of underwater speakers, showed responses from 100Hz-450Hz and 950Hz-1kHz, without finding responses to intermediate frequencies [20].

To avoid the complications that arise from the air-water interface and the interference generated by arrays of speakers, we attached a mini-speaker directly to the back coverslip wall of our 3D printed imaging chamber, effectively turning the coverslip into a water-coupled speaker (Figure 1A). The fidelity of the resulting stimuli was tested with a hydrophone, confirming the generation of the target frequencies without appreciable interference or harmonics (Figure S1).

**Figure 1.**
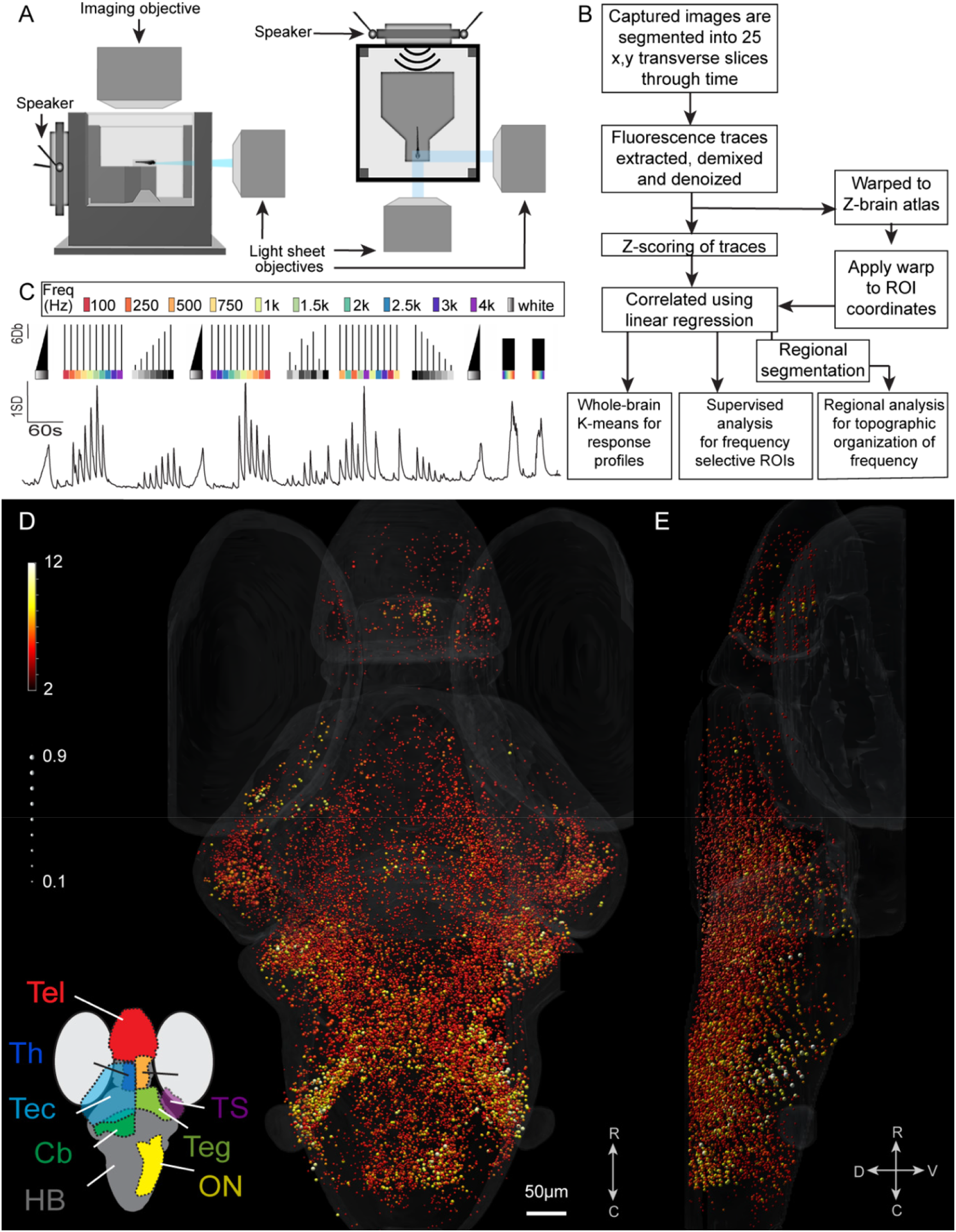
Experimental design, analytic workflow, and auditory-responsive ROIs. **(A)** Schematic of the imaging and experimental setup used to observe brain activity from zebrafish larvae, showing the mini-speaker attached to the back coverslip of the chamber. **(B)** Summary of the analysis workflow used to identify and categorize auditory neurons across the brain. **(C, top)** shows a schematic of the stimulus train used, including stimuli with a range of different forms, frequencies, and amplitudes. **(C, bottom)** shows the mean response of all the auditory-responsive ROIs in the resulting dataset. **(D and E)** All auditory responsive ROIs across the brain are shown with the strength of the response represented by the color, and the correlation to the linear regression model (r^2^ value) represented as the size of the sphere. The strength of the response is between 2 (the threshold for inclusion) and 12SD. Brain regions containing a high proportion of auditory-responsive ROIs are identified spatially **(Inset, D)**. Tel: telencephalon, Th: thalamus, Tec: tectum, TS: torus semicircularis, Teg: tegmentum, Cb: cerebellum, ON: octavolateralis nucleus, HB: remaining hindbrain. R, rostral; C, caudal; D, dorsal, V, ventral here and in subsequent figures. Scale bar applies to both **D** and **E**.

Using this approach, we ensonified the chamber with a diverse set of auditory stimuli (Figure 1C), which fell into two basic categories: pure tones of individual frequencies ranging from 100Hz to 4kHz, and white noise stimuli containing frequencies throughout this range. The pure tones were used to explore frequency selectivity, while white noise stimuli of various amplitudes allowed us to study response sensitivity. These stimuli lasted for 1 second without changes in frequency or amplitude. We also included white noise amplitude ramps, which were 20 second stimuli with an amplitude increasing from silence to full volume, and frequency sweeps with ascending or descending frequencies between 100Hz and 4kHz across a 30 second stimulus.

To observe brain-wide responses to these stimuli, we performed volumetric calcium imaging using dual plane selective plane illumination microscopy (SPIM, Figure 1A) and the *HuC:H2B-GCaMP6s* transgene line [30, 31]. Volumetric images were collected, processed, and analyzed using the neuroinformatic pipeline outlined in Figure 1B (see Vanwalleghem, et al. [32] and Methods for details). Briefly, we started by segmenting regions of interest (ROIs) generally corresponding to individual neurons, and then used linear regression and thresholding to identify all auditory-responsive ROIs (Figure 1B). The mean signal of these collective ROIs showed responses to all stimuli in our stimulus train except for pure tones at high frequencies (Fig. 1C). Using the 3-dimensional positions of each ROI, registered against the Z-brain atlas of the larval zebrafish brain [33](Figure 1B and Methods), we mapped all responsive ROIs onto a reference brain, including the strength (regression coefficient) and correlation coefficient of the auditory responses of each ROI (Figure 1D and E).

The resulting map of brain-wide auditory responses revealed ROIs throughout numerous regions of the brain, including all regions that have previously been described as auditory-responsive in larval zebrafish zebrafish [19, 20, 34, 35]. Auditory responses were particularly dense and strong in the octavolateralis nucleus (ON), which receives direct innervation from the ear [36, 37] and relays auditory information to other regions of the brain [38, 39]. We also observed robust auditory responsiveness in downstream structures including the torus semicircularis (TS), remaining hindbrain (HB), thalamus, and cerebellum. Responses were sparser, but nonetheless clear, in the tectum, telencephalon, tegmentum, and pretectum, with a small number of ROIs in the habenula. We also used an established method to detect stimulus-associated drops in the GCaMP signal that are indicative of inhibition ([40], see Methods), but found no consistent evidence of auditory-responsive neurons that were inhibited.

While these results reveal the locations of auditory neurons across the brain, they do not demonstrate whether different categories of neurons, with selective responses to particular stimuli, exist within this system. To explore this possibility, we performed k-means clustering (see Methods) to identify functional categories (clusters) of ROIs responding to individual sound properties or categories of properties. Using this approach, we found a total of six categories of response types (clusters). Two of these clusters responded to all the stimuli except for the highest frequency pure tones (Figure 2A, 2B). The first of these (Figure 2A) showed its strongest responses to white noise stimuli, including those at a low amplitude. The second broadly tuned cluster (Figure 2B) was less sensitive to white noise stimuli but showed stronger responses to pure tones across a range of frequencies. Interestingly, ROIs belonging to these clusters were restricted to the ON, and to a much lesser degree, the TS. Given that the ON, which is homologous to the cochlear nucleus in mammals, is the first brain structure to receive information from the ear, this suggests that the early stage of auditory processing in larval zebrafish predominantly involves broadly tuned neurons, consistent with findings from other fish species [37, 39]. However, as these clusters’ preferences for white noise or pure tones were not absolute, it is possible that there is a range of broadly responsive ROIs in the ON, and that these two clusters represent two halves of a diverse continuum of such neurons.

**Figure 2.**
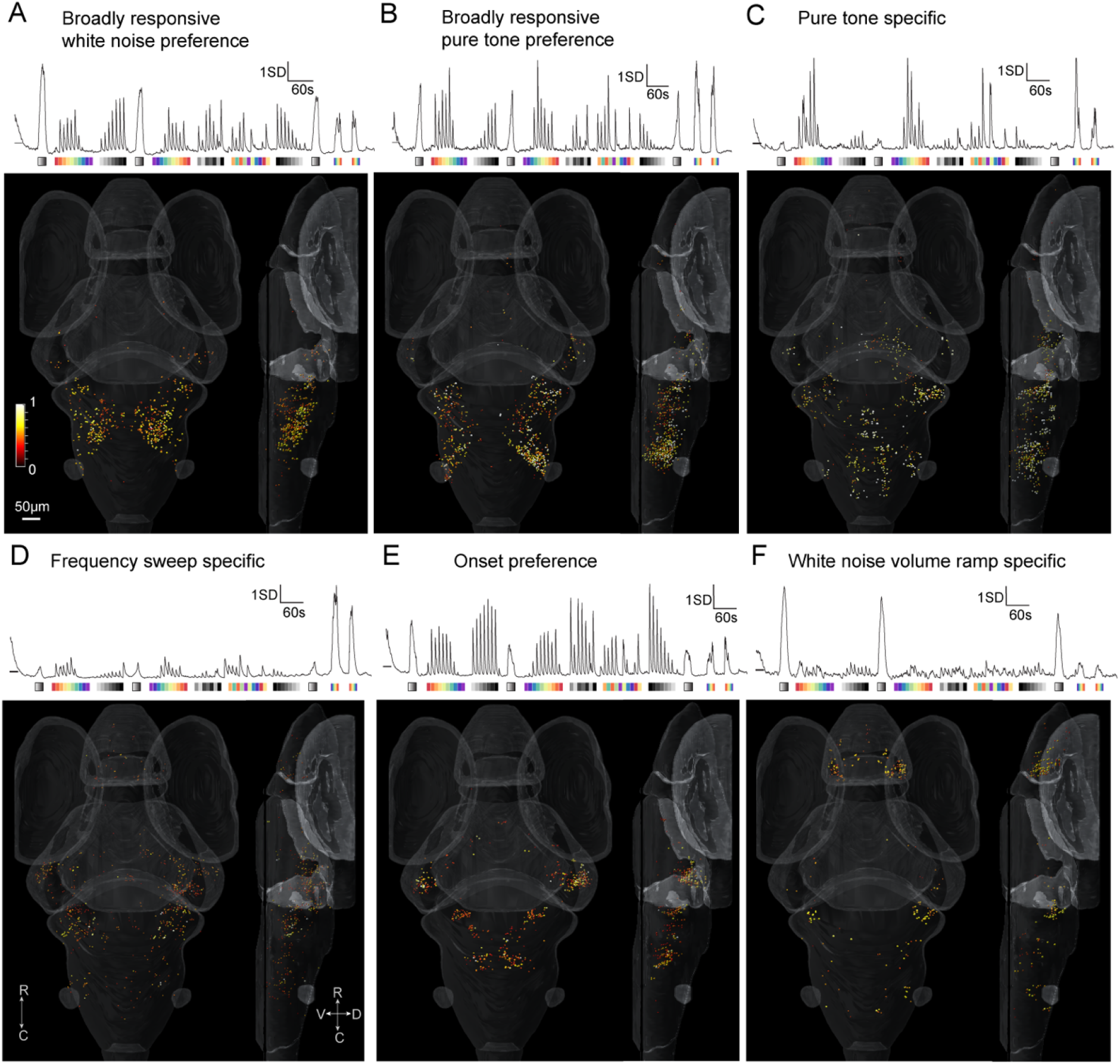
K-means clustering of brain-wide auditory responsive ROIs. K-means clustering revealed 6 clusters of ROIs that respond to particular properties of auditory stimuli. Broadly tuned clusters were detected **(A, B)** that were predominately located in the ON. A pure-tone specific cluster was present in the medial HB, lateral cerebellum, TS and pretectum **(C)**. The fourth cluster was responsive to the frequency sweeps, with ROIs in the TS, ON and lateral cerebellum **(D)**. The fifth cluster represents an onset cluster present in the TS, the lateral cerebellum and the medial HB **(E)**. The final cluster revealed that white noise volume ramp-selective ROIs are present in the pallial region of the telencephalon, the lateral cerebellum and sparsely in the HB **(F)**. Scale bar applies to all panels.

A third functional cluster included ROIs that responded almost exclusively to pure tones across a range of frequencies (Figure 2C). These ROIs were distributed across the ON, TS, HB, and tegmentum. This distribution closely resembles that of the fourth cluster (found in the ON, TS, tectum and remaining HB and sparsely in the thalamus and telencephalon, Figure 2D), in which the ROIs were specifically responsive to frequency sweeps. These clusters have responses to pure tones in common, whether those tones occurred in isolation (Figure 2C) or at different times during a frequency sweep (Figure 2D), suggesting that the auditory ROIs in these regions are involved in the detection of pure tones, and potentially the discrimination of different frequencies.

A fifth cluster contained ROIs that responded strongly to stimuli with sudden onsets (square tones), including both pure tones and white noise stimuli and stimuli with very low amplitudes. In contrast, they had relatively weak responses to auditory ramps, even once these ramps reached their peak amplitudes (Figure 2E). We interpret these to be “onset” ROIs that are specifically sensitive to the sudden occurrence of sound. These ROIs were particularly abundant in the TS, and were also present in the ON, and to a lesser degree, the thalamus. This cluster indicates that the larval zebrafish TS is particularly sensitive to the onset of a sound, spanning from the lowest frequency played (100Hz), through to 3kHz, and to all white noise volumes. A reciprocal cluster (Figure 2F) responded almost exclusively to white noise ramp stimuli, with little or no response to stimuli with sharp onsets. This cluster’s distribution was distinct from those of the other five clusters, with ROIs predominantly restricted to the lateral pallium and lateral cerebellum. Taken together, these findings indicate that larval zebrafish have neurons selective to a range of specific properties of sound by 6dpf.

The application of a clustering approach with our brain-wide imaging data provides a framework for understanding more nuanced auditory processing than has previously been described in any larval fish species. The identification of these diverse and specialized auditory neurons, with selective responses to pure versus complex tones, and to the onset of square tones versus ramps, suggests that some of the sophistication of the adult fish auditory system [3, 24, 37, 38, 41, 42] is already in place in zebrafish larvae at 6 dpf. There are also hints that the structure of this adult pathway is present, in a nascent form, in these larvae. Our results reveal that broadly responsive auditory neurons are concentrated in the ON, at the early stages of auditory processing, whereas neurons with more specialized responses are likely to be located downstream of the ON in the TS, or in a variety of brain structures located later in the auditory processing pathway. Even within brain regions, we found examples of spatial segregation of neurons with distinct response profiles. These include the separate regions of the ON occupied by neurons with preferences for white noise and pure tones (Figure 2A, B) and distinct distributions in the hindbrain for neurons responding to pure tones, onset stimulus, and white noise ramps (Figure 2C, E, and F). These distinct spatial distributions are difficult to interpret in larvae, where these structures have not yet nucleated, but they nonetheless suggest spatially heterogeneous encoding of the individual components of sound, likely representing the precursors of the mature auditory system.

Notably, we observed ROIs that were responsive to pure tones, raising the possibility of frequency discrimination in larval zebrafish. Adult zebrafish have been shown to discriminate different frequencies in behavioral experiments [43], but it is less clear whether such discrimination takes place in any larval fish (zebrafish or other species). Our previous work has suggested that there is little or no frequency discrimination among ON neurons in larval zebrafish [19], although a recent study from Privat, et al. [20] proposed distinct neural pathways for broad bands of high and low frequency sounds at the whole-brain level. Overall, the larval zebrafish literature suggests a detection range up to roughly 1kHz, with, at best, coarse frequency discrimination across this range [7, 25, 29]. Having identified a large population of ROIs across the brain that respond to pure tones (Figure 2), we next explored whether their responses provide evidence for frequency discrimination or the topographic representation of different frequencies across or within brain regions.

To conduct this analysis, we adopted a supervised approach, building regressors for each frequency of pure tone in our stimulus train. We then defined an ROI as frequency selective if its regression coefficient (response strength) was greater than 2.5SD above the mean distribution for that frequency, and below 2.0SD for all other frequencies (see Methods). Applying these criteria to 35,720 auditory-responsive ROIs across seven larvae, we identified 9,411 ROIs that were frequency selective, including ROIs selective to each of our 10 frequencies ranging from 100Hz to 4kHz (Figure 3A). The average traces of neurons selective to each frequency peaked much higher (>two-fold) than adjacent frequencies through the low-mid frequency range (up to 2kHz) (Figure 3A). In contrast, those for 2.5k, 3k, and 4.0kHz-responsive ROIs were similar to the adjacent frequencies (Figure 3A). This suggests that larval zebrafish may be sensitive to these higher 2.5-4kHz frequencies without being able to discriminate among them, whereas for the lower frequencies, discrimination appears likely. We then visualized the frequency-selective ROI locations throughout the brain (Figure 3B, C). Although we found most of the ROIs spanned the rostro-caudal extent of the brain, the most responsive ROIs were located in the hindbrain, including the ON. This was especially true for ROIs that were selective to the mid frequencies (750Hz-2000Hz). The low (100Hz) and high (2.5-4.0kHz) frequencies were more broadly dispersed along the rostro-caudal axis, including notable concentrations of ROIs in the telencephalon (Figure 3D). For frequencies above 2.5kHz, these responses tended to involve the lateral telencephalon, where ROIs responsive to lower frequencies were concentrated in the medial telencephalon (Figure 3B).

**Figure 3.**
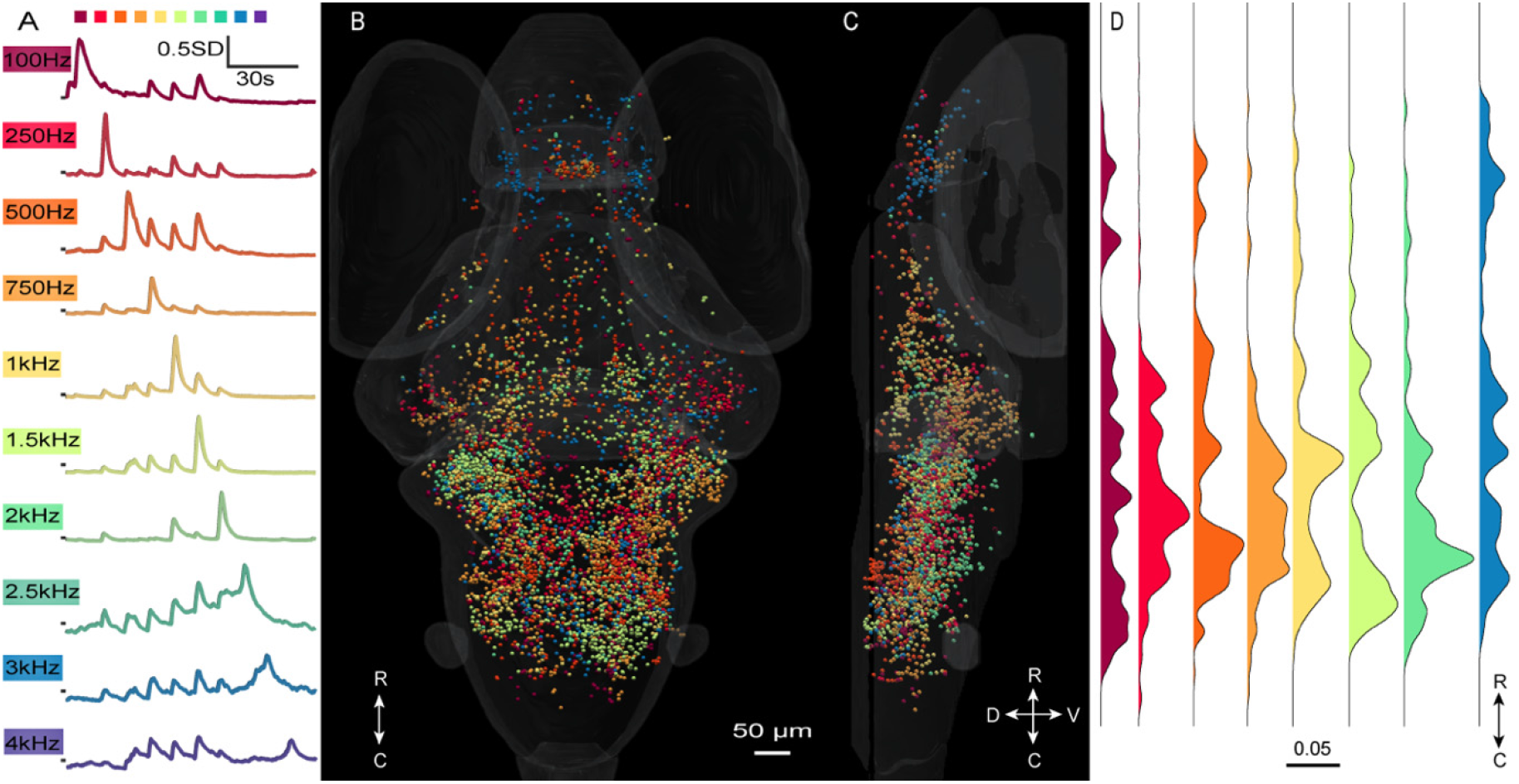
Frequency specificity and homogeneity at the whole-brain level. **(A)** shows the mean responses of all ROIs selectively responding to each frequency in our stimulus train **(top**), and **(B)** shows their positions in the brain from a dorsal view, and **(C)** shows a lateral view. Scale bar applies to both **B** and **C**. **(D)** Using a one-dimensional kernel density estimation (KDE) computation, we looked at the density of frequency-selective ROIs at the whole-brain level, from rostral to caudal. Whole-brain images such as those in **(B)** can be seen for individual frequencies in Supplementary Figure 2.

In mammals and some adult fish, auditory responses are organized tonotopically, whereby the frequency specificities of the neurons correlate with their spatial positions [38, 42, 44–46]. In larval zebrafish, one study has hinted at tonotopy, albeit inconclusively [19], whereas another has suggested broad brain-wide segregation of high and low frequency responses without responses to intermediate frequencies [20]. To search for topographical encoding of frequency in our dataset, we carried out a spatial analysis of frequency selective ROIs in ten brain regions that were prominent in our brain-wide dataset or which are known to be involved in auditory processing [19, 20, 34, 35, 39]. These regions were the ON, TS, thalamus, pretectum, tectum, habenula, tegmentum, telencephalon, cerebellum, and remaining HB. Collectively, these regions include more than 80% of the ROIs that we segmented across the brain, and more than 97% of frequency-responsive ROIs in our dataset. Because of the overlapping responses and similar spatial distributions of ROIs responding to 2.5k, 3k, and 4 kHz tones (Figure 3), we combined these into a single category for this analysis.

The broad frequency tuning of each brain region can be gauged by comparing the abundance of the frequency selectivities of its ROIs to those of ROIs across the entire brain (Figure 4A). The most striking departures from brain-wide averages were found in the telencephalon, and to a lesser degree in the habenula, both of which were enriched for responses to high frequencies (Figure 3B). The thalamus was notable for lacking responses to both high and low frequencies, responding instead to frequencies from 500Hz to 2kHz. Other brain regions exhibited frequency-responsive ROIs whose relative proportions approximately reflected those across the brain as a whole.

**Figure 4 –.**
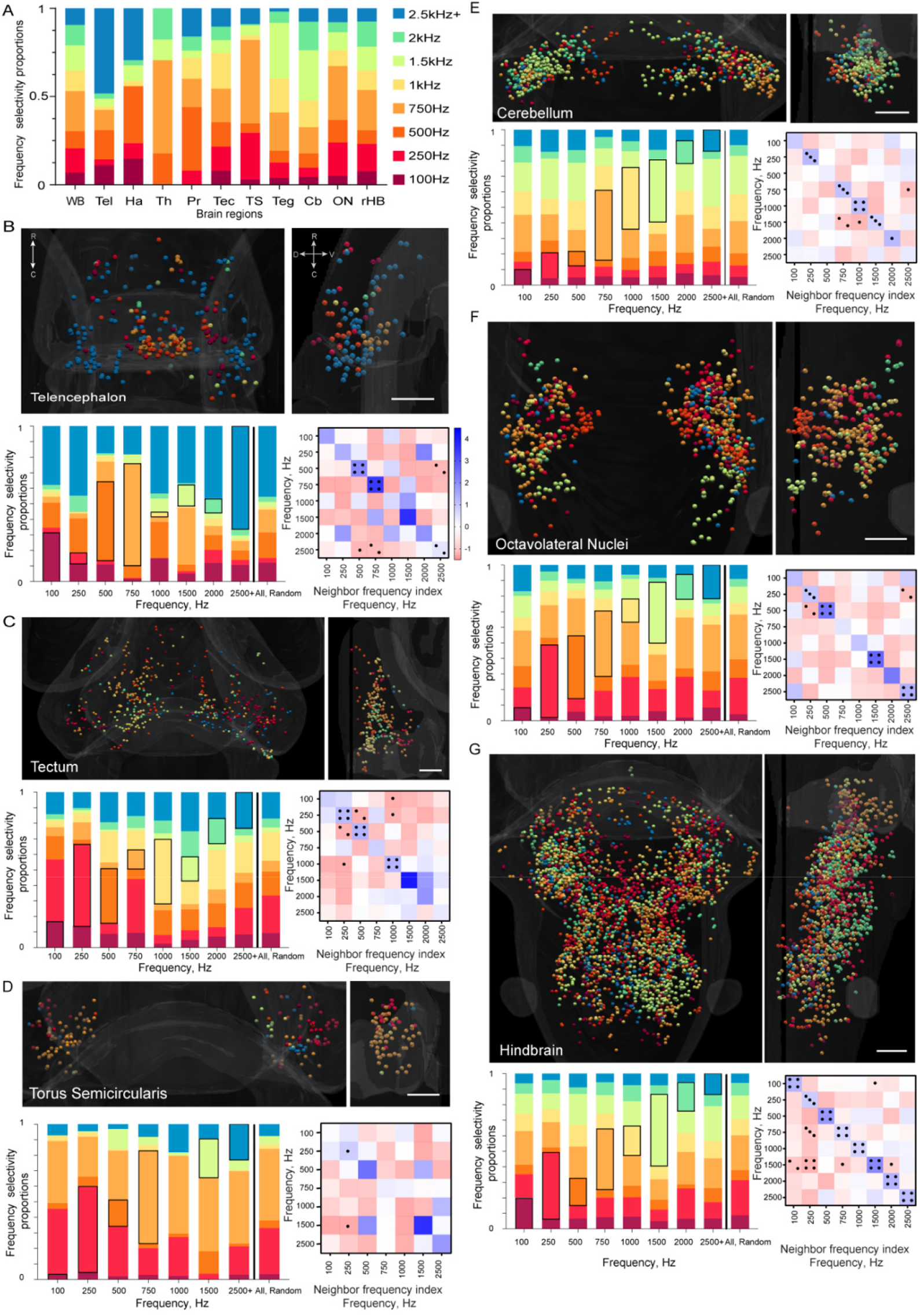
Regional frequency distribution and tonotopy –. **(A)** shows the fraction of ROIs per frequency at the whole-brain (WB) and regional levels. For **(B-G)**, the spatial distribution of ROIs from the dorsal **(top left)** and lateral **(top right)** views are shown. Bar graphs **(bottom left, B-G)** show the proportions of nearest neighbors for ROIs of a given frequency preference (separate bars for ROIs responding to different frequencies), with same-frequency neighbors indicated by a black box. For comparison, data from the spatially randomized dataset are shown in the last bar. Matrices **(bottom right)** of the neighbor frequency index show pairs of frequencies that are overrepresented **(blue, bar in B)** or underrepresented **(red)** as neighbors. Significant results versus the randomized dataset are indicated (P value: • <0.05, •• <0.01, •••<0.001, ••••<0.0001). Kruskal-Wallis tests with Dunn’s correction for multiple comparisons were performed to establish P-values. The k-nearest neighbor heat map legend is shown in **(B)** and applies to panels **(B-G)**. The same information for the remaining brain regions can be found in Supplementary Figure 3 and all P-values are presented in Supplementary Figure 4.

To search for topography in each of these brain regions, we next performed a k-nearest neighbor (kNN) analysis whereby, for each ROI selective to a given frequency, we observed a selected number (*k*) of nearest neighbors, and recorded the frequency selectivity of those neighbors (see Methods). The results of this nearest neighbor analysis were then compared to a randomized dataset where the same number of ROIs of each frequency was retained, but the frequency selectivities of these ROIs were reassigned randomly. We performed 100 iterations of this randomization, and compared the neighbors yielded from these random datasets to those seen in our actual imaging dataset (Figure 4). The rationale for this approach was that heterogeneous distributions of frequency-selective neurons, where neurons responding to a particular frequency are spatially clustered, should result in a higher prevalence of same-frequency neighbors in our experimental dataset versus the randomized datasets. Further, if frequency responsiveness is spatially organized in a tonotopic map, ROIs responding to adjacent frequencies should be overrepresented among the neighbors in the experimental versus the random dataset.

Each brain region had its own profile of frequency-responsive ROIs (Figure 4A), and in each brain region, the randomized nearest neighbor population closely reflected this overall abundance of each frequency (Figure 4B-H, “all random” bar, *k* neighbors listed in Methods). This provided assurance that our randomization was working. We could identify spatial clustering of the ROIs of each frequency by looking for cases where the same frequency was overrepresented among its neighbors (dark boxes in each bar), as compared to neighbors from the “all randomized” bar. To quantify these effects, we generated a “neighbor frequency index” (see Methods). This gives positive values (shown in blue, Figure 4B-G) when a given frequency is overrepresented among the neighbors and negative values (red) when that frequency is underrepresented. Accordingly, spatial clustering of ROIs with the same frequency sensitivity should be represented by positive values along the diagonal of each matrix (bottom right, Figure 4B-G), and tonotopy would drive positive values in squares adjacent to the diagonal.

In the thalamus, the pretectum, and the habenula, there were not enough ROIs to support a statistical analysis of their spatial distribution (these distributions are shown in Supplementary Figure 3). In the tegmentum, there were no statistically significant departures from a homogeneous distribution of ROIs responding to different frequencies, suggesting a lack of spatial frequency representation (Supplementary Figure 3). Within the telencephalon, tectum, TS, cerebellum, ON, and HB, however, there were pronounced relationships between the spatial distributions if ROIs and their frequency preferences (Fig 4B-G).

Using our kNN analysis, we found numerous cases in which same-frequency ROIs were spatially clustered in other auditory brain regions (Figure 4B-G). In most regions, there were examples of significantly elevated neighbor frequency indices for ROIs responding to the same frequency (blue diagonal lines, Figure 4B-G), with these being particularly prominent in the tectum, cerebellum, ON, and hindbrain, where all same-frequency indices were elevated, many of them significantly. This was not consistently the case in the TS, where the very high abundance of 250 and 750Hz-responsive ROIs may have limited the sensitivity of the frequency neighbor index. The telencephalon is notable for its segregation of high and low frequency ROIs. Low frequency responses are abundant in the medial telencephalon, which is homologous to the amygdala in mammals [47, 48],whereas high frequency responses are concentrated in the lateral telencephalon, which corresponds to the mammalian hippocampus [49, 50].

Although our nearest neighbor analysis identified numerous examples of spatially clustered same-frequency ROIs, it provided scant evidence for tonotopy, in which responses to similar frequencies would be spatially adjacent. The distributions of ROIs in the primary auditory region, the ON (Figure 4F), appeared to have the lower frequencies spatially organized in the dorsomedial section, with the populations representing the higher frequencies being located more caudally. The tectum (Figure 4C) also appeared to contain a medial (high frequency) to lateral (low frequency) gradient, while the HB (Figures 3D and 4G) appeared to have a rostral (low frequency) to caudal (middle frequency) gradient. None of these trends, however, was confirmed by consistently elevated frequency neighbor indices for consecutive frequencies, which would have appeared as significantly blue cells adjacent to the diagonals in Figure 4. Such tonotopy is a hallmark of the mammalian cochlear ganglion [44, 51, 52], and is also present in the ON and TS in the adults of other teleost fish species [38, 42, 53, 54], but our results suggest that this tonotopy is not present in zebrafish at 6dpf. One possibility is that tonotopy may develop at later stages of development in zebrafish. Another possibility is that zebrafish may have no tonotopic organization for sound frequencies, perhaps because spatial organization is dependent on the cochlea, which is absent in teleosts.

Overall, the observations from our nearest neighbor analysis suggest that, while neurons responding to a given frequency are spatially clustered at multiple stages of the auditory processing pathway in larval zebrafish, systematic tonotopy is absent or still in a rudimentary form. Details of our approach and statistical tests mean that these represent conservative interpretations. We observed several cases where strong positive neighbor frequency index scores proved to be non-significant. These generally occurred where the brain region as a whole (the TS, for instance) or the frequency being tested (1500Hz in the tectum, for example) had a small number of ROIs. It is likely that larger datasets would reveal significance in these relationships, as they did in the HB (Figure 4G) where a large number of ROIs show significance in all same-frequency comparisons. We also note that our cases of strong same-frequency clustering may be masking tonotopy, given that a high abundance of same-frequency ROIs in a neighbor pool depletes the pool for other frequencies, including adjacent ones. This may provide a partial explanation for the lack of significant tonotopy in our analyses, although it is also possible that this tonotopy simply does not exist in zebrafish larvae.

These results differ in important ways from two past studies of brain-wide auditory processing in larval zebrafish. One of these, from our own group [19], showed a limited frequency response range (from 100-400Hz), and fewer responsive brain regions. In large part, this is likely due to a stimulus train that was limited to a 100-800Hz range, and that comprised only pure tone bursts. Furthermore, we used an air-coupled speaker in these past experiments, raising the likelihood that the air-water interface interfered with the delivery of our auditory stimuli. Finally, this study was done on a more primitive light-sheet microscope, which likely limited our detection of weak signals and deeper brain regions.

A recent study by Privat, et al. [20] showed two regions of frequency sensitivity, ranging from 150-450Hz and 950Hz-1kHz (the highest frequency tested), which were represented primarily in the hindbrain and midbrain, respectively. No responses were observed to frequencies between these ranges. Their use of geometric ROIs presumed to encompass numerous neurons could explain the absence of intermediate frequencies from their study, consistent with the fact that we often found our 500Hz- and 750Hz-responsive neurons intercalated with ROIs responding to other frequencies. Furthermore, they used a cytoplasmically-targeted GCaMP, providing the benefit that they could analyze responses in neuropil region (which our nuclear-targeted GCaMP did not reveal), but at the cost of resolving individual neurons. Finally, their study targeted particular parts of the brain for imaging and may therefore have missed responses that we have found in regions not previously associated with auditory processing in this system.

Previous studies have described the auditory sensitivity and hearing range of larval zebrafish using a range of different approaches [5, 7, 19, 20, 25], generally describing a system with limited sensitivity to stimulus components, a tight frequency range, and little spatial specialization in auditory coding. In this study, we have used a more diverse stimulus train, a novel method for sound delivery, and an upgraded imaging system to detect and characterize the brain-wide responses of individual neurons in larval zebrafish. We have revealed response characteristics that suggest broad auditory tuning in the early stages of the network, especially in the ON, and sensitivity to more nuanced stimulus properties in later stages, including in the TS, telencephalon, cerebellum, and remaining HB. These later steps may provide sensitivity to pure tones, the detection of the onset of auditory stimuli, and the detection of stimuli that ascend in intensity gradually. We have also shown broader frequency sensitivity, up to 4kHz, than has previously been observed, and spatial clustering of frequency-selective neurons in several brain regions, although does not reflect coherent patterns of tonotopy.

These results provide evidence for more sophisticated auditory processing than has previously been appreciated in this model system and offers a starting point from which to use its strengths in relation to imaging and optogenetics to characterize the circuits and networks composing the auditory system as a whole.

## Acknowledgements

We thank the University of Queensland’s Biological Resources aquatics team for animal care. We also thank Emmanuel Marquez-Legorreta for his intellectual expertise regarding telencephalic activity. We thank Germán Sumbre and Rowan Tweedale for suggestions on the manuscript. Support was provided by an NHMRC Project Grant (APP1066887), a Simons Foundation Pilot Award (399432), a Simons Foundation Research Award (625793), and two ARC Discovery Project Grants (DP140102036 & DP110103612) to E.K.S.; an EMBO LongTerm Fellowship to G.C.V.; a fellowship from the Human Frontier Science Program (LT000146/2016)to M.A.T.; and University of Queensland Postgraduate Awards to R.E.P and L.S..

## Author Contributions

Conceptualization, E.K.S.; Methodology, R.E.P.,G.C.V., I.A.F-B and L.S.; Investigation, R.E.P., G.C.V.; Animal Colony Maintenance, L.C.; Formal Analysis, R.E.P.,_G.C.V. and L.S.; Data Curation, R.E.P, G.C.V..; Writing–Original Draft, R.P. and E.K.S.; Writing–Review and Editing, G.C.V., L.S., L.C. and I.A.F-B.; Figure Construction, R.E.P. and L.S.; Funding Acquisition, E.K.S.; Resources, E.K.S.; Supervision, E.K.S. and L.C.

## Data Availability

The dataset generated and analyzed for this study can be found in the UQ eSpace https://doi.org/10.14264/680dfce, and the code used in the analysis can be found at https://github.com/Scott-Lab-QBI/Brainwide_auditory_processing.

## METHODS

### Animals

Adult zebrafish (*Danio rerio*) were maintained at 28.5°C at a density of 10-15 fish per liter and on a 14/10 hour light/dark cycle. The *HuC:H2B–Gcamp6s* transgenic line was used for these experiments, targeting the calcium indicator GCaMP6s to the nuclei of all neurons[30]. Fertilized eggs of the TLN strain were transferred into E3 medium (distilled water with 10% Hanks solution, consisting of 137mM NaCl, 5.4mM KCl, 0.25mM Na_2_HPO_4_, 0.44mM KH_2_PO_4_, 1.3mM CaCl_2_, 1.0mM 654 MgSO_4_ and 4.2mM NaHCO_3_ at pH 7.2) and kept at 28.5°C in an incubator with a 14/10 hour light/dark cycle. Zebrafish housing, breeding, larval maintenance, and experiments were performed with approval from the University of Queensland Animal Ethics Committee (IMB/237/16/BREED and SBMS/378/16).

### Calcium imaging

6dpf larval zebrafish were set in a 2% low melting point agarose (Progen Biosciences), dorsal side up, and placed it into the custom-made chamber [35]. The 3D printed 24×24mm chamber consists of a square plastic base and a 0.2mm^2^ post in each corner. 20mm x 20mm glass coverslips (ProSciTech) were fixed on to each side using a waterproof glue (Liquid Fusion Clear Urethane Adhesive). The glass coverslips enabled light-sheet illumination from the front and one side of the animal with minimal light distortions. Audio stimuli were delivered from the speaker adhered to the rear coverslip. Agarose-set fish were mounted onto the platform of the chamber with additional agarose to prevent the fish from moving throughout the experiment. Once the agarose had set, the chamber was filled with E3 medium and allowed to sit for a minimum of 30 minutes to minimize drifting of the fish during imaging.

Whole-brain calcium fluorescence imaging was done using a custom-built selective plane illumination microscope (SPIM) to determine the neural responses *in vivo* while the auditory stimuli were presented [31, 55]. The fish was simultaneously illuminated with two planes from the front and one side and imaged at 10μm increments in the dorsoventral axis with an exposure time of 10ms. This produced a 25-slice volumetric representation of the entire brain, with a 4Hz volumetric imaging rate. Details of this microscope and imaging procedure have been described previously [17, 31].

### Auditory stimulation

Auditory stimulation was provided by a mini speaker (Dayton Audio DAEX-9-4SM Skinny Mini Exciter Audio, Haptic Item Number 295-256) fixed to the back glass surface (caudal to the animal) of the imaging chamber and wired to an amplifier (Dayton Audio DA30 2 x 15W Class D Bridgeable Mini Amplifier). Preliminary testing was done to determine the optimal speaker capabilities (data not shown). The sound pressure level (SPL) reading was taken before each experiment in each chamber. Sound level measurements of the white noise at a playback volume of 0dB digital full scale (FS) were taken at the position of the fish before filling the chamber and measured approximately 84dB (SPL) noting the background noise was 40-45dB (SPL). A measurement of 74-76dB (SPL) was taken at the surface of the liquid when the chamber contained E3 medium. This ensured consistency between chambers and between experiments.

Stimulus playback and image acquisition were performed using Micro Manager software [17, 56]. Each presentation of the stimulus train included a 20 second white noise amplitude ramp, 1 second of 10 frequencies with a 2ms rise and fall time (100, 250, 500, 750, 1000, 1500, 2000, 2500, 3000, 4000Hz) at 0dBFS with 9-second inter-stimulus intervals (ISIs), and one second white noise with a 2ms rise and fall time at 7 amplitudes ranging from −18dBFS to 0dBFS in 3dB increments with a 9-second ISI. This train was presented 3 times, changing the order of the type of stimulus, and using ascending, descending and quasi-random orders of frequencies, and ascending, quasi-random and descending orders of amplitudes. There was 30-second rest between each stimulus type and stimulus presentation.

To test the frequency response of the speaker, a custom-built hydrophone (Neptune Sonar, UK) was used to take spectral analysis recordings of the white noise and individual frequencies while the chamber contained E3 medium, and without the fish present. The recording was done by placing the microphone in the water-filled chamber. This was then connected to an Edirol FA-66 microphone preamplifier audio interface (Roland, Japan) and recorded into a professional audio program (Ableton Live!, Germany). The spectral analysis was performed using a Fast Fourier Transform plugin SPAN (Voxengo, Sweden) and showed the white noise response at lower frequencies attenuating at 150Hz, and high frequencies beginning to attenuate at 6kHz. The recordings also delineated one of the limitations of the speaker delivery system design. There was a comparably low frequency response at 100Hz in the white noise and individual frequency recordings. The individual pure tones contained minimal harmonic distortion. The frequency showing the most secondary harmonics was 250Hz. This was approximately 21dB below the fundamental, meaning that the fundamental was over three times the amplitude of any harmonic across all frequencies, which was at or below the hearing threshold of the fish [35]. Secondary harmonics for other frequencies were all more than 40dB below the fundamental. During the experiments, the chamber was fastened to the platform to minimize any sonic or motion artefacts that might have resulted from vibrations.

### Data processing and analysis

We excluded three fish because the imaging quality was blurry, or the fish were tilted. Of the remaining ten fish, once the images were captured, videos were cropped, the transverse slices were segmented to identify ROIs corresponding to individual neurons, and the data were resaved as 25 individual Z-stacks per experiment over time, using Image J v1.52c. Each of the 25 planes was then motion corrected using the NoRMCorre algorithm [57]. Fluorescent traces generated by calcium transients in each ROI were then extracted, demixed, and denoised using the CaImAn package previously described [17, 19, 57]. These traces were then z-scored, and correlated using linear regression, which was built from the stimulus train.

MATLAB v9.5 (Mathworks) was used to further analyze the data. Linear regression was used to extract the auditory-responsive ROIs, and a regressor was built for each of the stimulus types: white noise ramps, individual pure tone sine waves, short white noise volumes, and frequency sweeps. This gives an indication of baseline and stimulus-driven activity and identified neurons that respond to the stimulus train. The motion-corrected 3D volumes were registered, using Advanced Normalization Tools (ANTs, https://github.com/ANTsX/ANTs), to the H2B-RFP reference of Zbrain [33, 58]. The resulting warps were then applied to the centroid positions of the ROIs, which allowed us to use their location within brain regions outlined by Zbrain to conduct region-specific analysis [32]. Three additional fish did not successfully warp to Zbrain and were excluded at this point, leaving an n=7.

Response types to the properties of sound were then categorized at a brain-wide level. We looked at the correlation coefficient of the fluorescence traces to each auditory stimulus type (10 pure tones, white noise volumes, frequency sweep for both increasing and decreasing frequency order), thresholding at an *r^2^* value of 0.1 and a regressor coefficient (response strength) threshold of 2.0 SD above the mean response. K-means clustering was then used as part of the analysis to look at brain-wide profile response types. Given that k-means forces every ROI into a cluster, we used an additional filter of an *r^2^* value of 0.5, when compared to the mean of that cluster to further clean up the data and remove ROIs that did not match the cluster. The analysis returned 15 clusters. From this, 3 non-auditory clusters, 4 clusters that were under-represented in more than half the fish and 2 noisy clusters were excluded, with the remaining 6 clusters were kept.

To look for frequency heterogeneity, a supervised conditional analysis was conducted to determine whether any of the ROIs were responsive to an individual frequency. For an ROI to be characterized as being frequency specific, its response must be 2.5SD above the mean response strength for that particular frequency and its response must be less than 2SD above the mean response strength at all other frequencies. We looked at responses at thresholds from 1.0 SD to 3.0 SD and determined that 2.5 SD provided the most accurate representation of the data, as 1SD left too many noisy responses, and 3.0 SD left very few ROIs passing threshold. We first looked at this across the brain and subsequently in all the auditory regions to determine if there were whole-brain [20] or regional [19] spatial representations of frequency. Regionally, we looked at areas previously indicated to be responsive to sensory processing, namely, the ON, TS, thalamus, telencephalon, cerebellum, tectum, tegmentum, habenula and HB. We also used the inverse criteria to look for potential ROIs whose response would be inhibited at a specific frequency (response had to be 2.5SD below the mean response strength to a particular frequency), but identified no inhibited ROIs [40].

### Data visualization

To give a visual representation of the data, clusters were mapped back onto the brain using Unity™ which has been adapted into a data visualization system. An isosurface mesh of the zebrafish brain was generated from the Zbrain masks for the diencephalon, mesencephalon, rhombencephalon, telencephalon and eyes using ImageVis3D [17, 18]. The mesh was imported in Unity and overlaid to the ROIs. Each ROI was represented as a sphere within the brain. This enabled a 3D visualization of the resulting clusters and provided a qualitative way to determine if there were any spatially significant results.

### Statistical analysis

To quantify spatial frequency heterogeneity at a brain-wide and regional level, a multi-dimensional density estimation was done to determine how the individual frequencies were arranged spatially at a whole-brain level (Figure 3D). This calculated the density of each frequency on the rostro-caudal axis. In order to characterize whether the frequency-selective ROIs showed any kind of spatial segregation, a k-Nearest Neighbor (kNN) analysis was carried out. This was done brain-wide and regionally. All code related to these computations is available in https://github.com/Scott-Lab-QBI/Brainwide_auditory_processing.

The spatial position of each group of frequency-selective ROIs was used as input to a 1D (Figure 3D) kernel density estimation (KDE) computation. The sklearn.neighbors.KernelDensity method from python’s scikit-learn module was used for this purpose. To compute the KDE, a Gaussian kernel was used and the optimal bandwidth was found through optimization of the log probability density under the KDE model with the sklearn.model_selection.GridSearchCV method. The range of bandwidth values was chosen to ensure that it covered three orders of magnitude up to a third of the maximum range of values in the dimension where the KDE was computed (e.g. if values in the rostro-caudal axis had a range of 1000 pixels, the range of bandwidths tested was from 3 to 300 pixels). For the brain-wide density plots in Figure 3D, the bandwidth was defined as a weighted average of the optimal bandwidths found for each frequency.

The characterization of topographic organization of frequency selective ROIs was performed through the kNN analysis. The kNN classifier receives a set of data points (the *xyz* coordinates of the ROIs in this case) and labels (the frequency for which the ROI is selective) and a parameter *k* and outputs a decision boundary in the *xyz* space that defines the most likely label for an ROI with those particular coordinates based on the labels of its *k* neighbors. In cases where a brain region contained a sufficient number of ROIs for each frequency (minimum of 4) in the right lateral area, only the ROIs that side were used. However, in two of the regions, the telencephalon and tegmentum, the ROIs located in the left lateral area were translated to be merged to the other area based on the detected midline (coordinate origin value of the medial-lateral axis, *y* = 315 pixels), with the merged set of points being used as the input.

The sklearn.neighbors.KNeighborsClassifier method was used together with sklearn.model_selection.cross_val_score in order to evaluate the accuracy of the classifier for values of *k* ranging from 1 to 100. Once the accuracy values were obtained, a *k* value was chosen so that it was not lower than 10 or higher than 20% of the total number of ROIs in that region. This means that we did not necessarily choose the k values that resulted in the highest accuracy but used the results to guide our choice of *k* to calculate the fraction of neighbors for each frequency (stacked bar graphs in Figure 4). The *K* chosen for each brain region was cerebellum: 20, hindbrain: 27, tectum: 13, ON: 19, TS: 11, telencephalon: 12, tegmentum: 27.

Finally, in order to quantify how spatially segregated the frequency-selective ROIs in each brain region were, a new set of ROIs was generated. The number of labels for each frequency was kept the same as for the raw data, but they were reassigned to different *xyz* coordinates using a uniform random distribution. The neighbors were then computed using the same *k* used in the real dataset. This was performed multiple times for each frequency with a different number of ROIs with new random label assignments (10, 50 and a maximum of 100 iterations). This analysis allowed us to identify how different the spatial distributions of the ROIs were when compared to the same set of xyz coordinates, but with frequency labels assigned randomly in space. The stacked bar graphs in Figure 4B-G show the mean of the neighbor fractions with all frequencies considered (e.g. the value of 100 Hz neighbors in telencephalon is the mean 100Hz neighbor fraction of the ROIs of all frequencies). This was done because the mean and standard deviation of the neighbor fractions across frequencies did not change with frequency. The heat maps show the ratio of the difference between the fraction of neighbors of each frequency in the raw data and the ROIs with randomized frequency labels which we defined as the neighboring frequency index (NFI). The NFI is obtained by *NFI* = (*n_real,freq_* – *n_random,freq_*) / *n_random,freq_*, where *n_real,freq_* is the neighbor fraction of the experimental data for a certain frequency and *n_real,random_* the neighbor fraction of the same frequency in a dataset containing the same numbers of frequency-responsive neurons of each type, but randomly reassigned to the ROIs in the dataset. The statistical analyses were done to compare the distribution of neighbor fractions of each ROI in the raw data per frequency and per brain region (*n* available in Supplementary Figure 2G) against the neighbor fractions of the 100 ROIs with random reassigned labels per frequency and brain region. We used the non-parametric Kruskal-Wallis test with Dunn’s correction for multiple comparisons to obtain the adjusted *P*-values.

