## Supplementary material for "Broad frequency sensitivity and complex neural coding in the larval zebrafish auditory system": Supplimental Figures 1-4

### Supplementary Figures

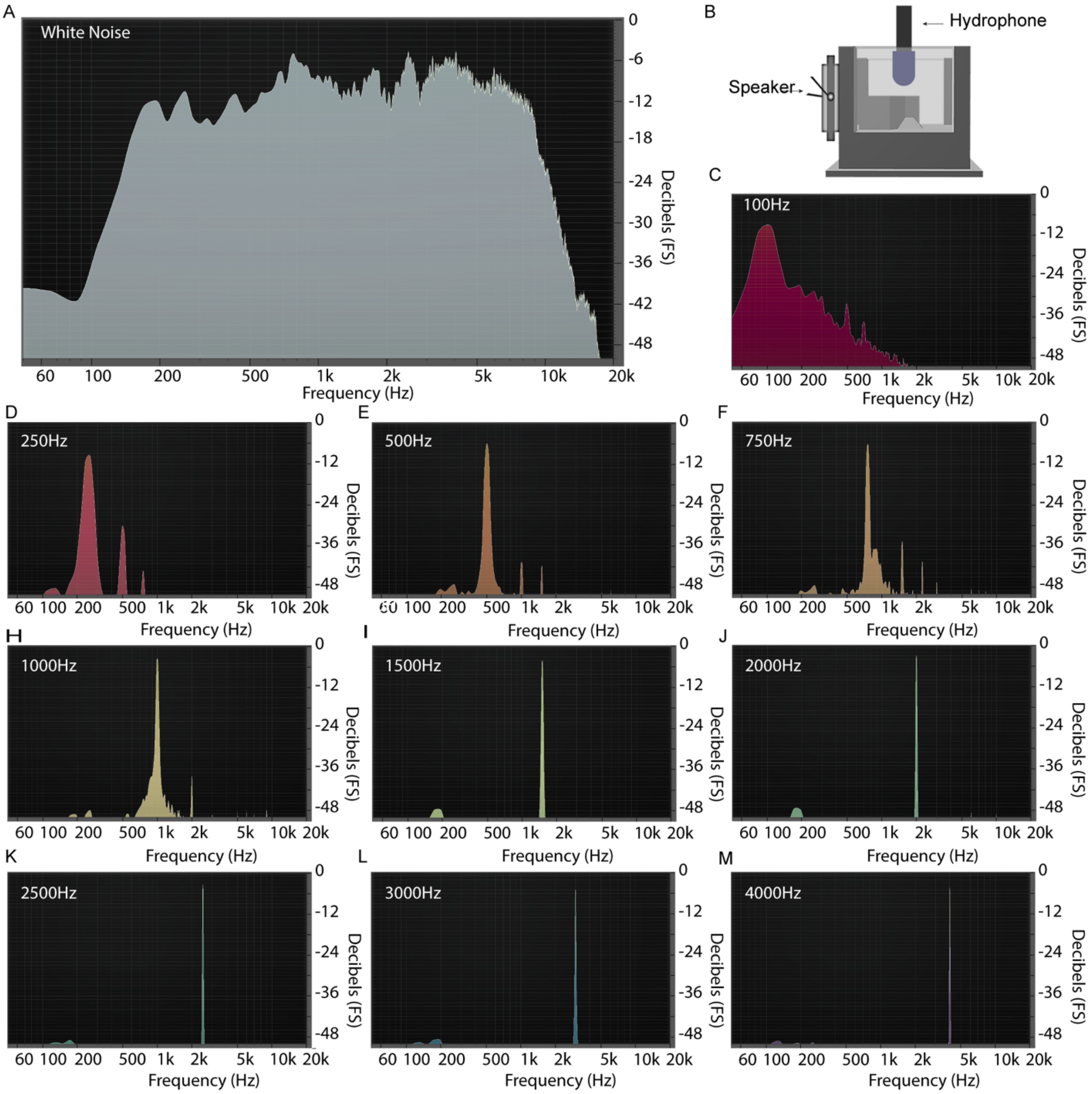


**Supplementary Figure 1 – Hydrophone recordings of white noise and sine wave stimuli within the chamber** – hydrophone recordings were taken using the custom built hydrophone (B) inside the water-filled chamber during white noise (A), and for each of the frequencies used in the stimulus train (C-M) of 100, 250, 500, 750, 1000, 1500, 2000, 2500, 3000 and 4000Hz (See Methods).

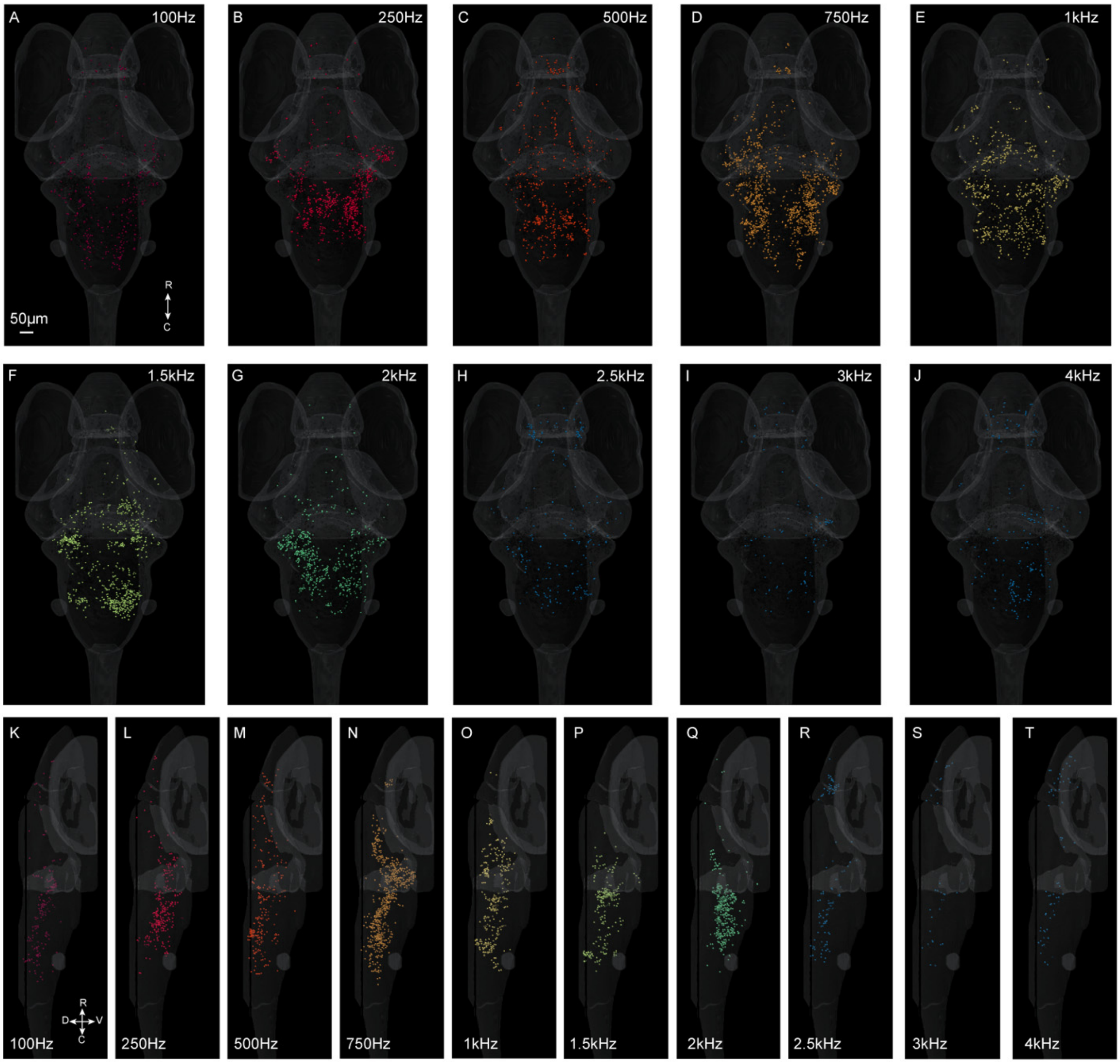


**Supplementary Figure 2 – Whole-brain images of individual frequencies** – dorsal (A-J) and lateral views (K-T) showing ROIs of each frequency used in the stimulus train that passed the supervised analysis thresholding. The relevant frequency is indicated in each panel.

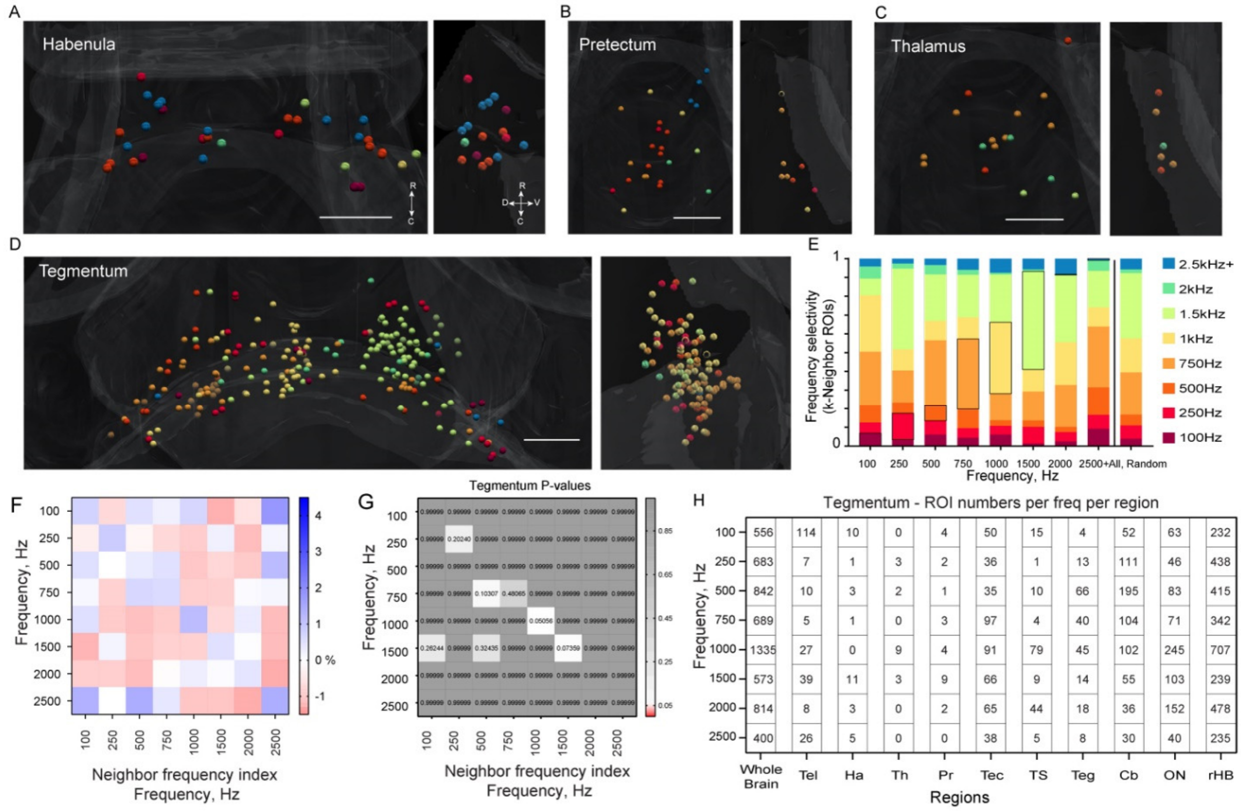


**Supplementary Figure 3 – Additional distributions of frequency sensitive ROIs**
(A-D) Unity images of the brain regions not shown in Figure 4 – the habenula (A), pretectum (B), thalamus (C), and tegmentum (D). Nearest neighbor analysis of the tegmentum is shown: bar graphs (E) show the nearest neighbors from the randomized dataset. The nearest neighbor value *k* varies per brain region (see Methods) for each frequency, from the real data and overall nearest neighbor fractions from the randomized dataset. Populations of same-frequency nearest neighbors are indicated with a dark outline. Matrices (F) of the neighbor frequency index show pairs of frequencies that are overrepresented (blue, bar in F) or underrepresented (red) as neighbors. P-values of the nearest neighbor analysis of the tectum (G, bar indicating color code), showing no significant separation between values from real and randomized datasets. (H) Total number of ROIs per frequency per brain region after the supervised analysis thresholding.

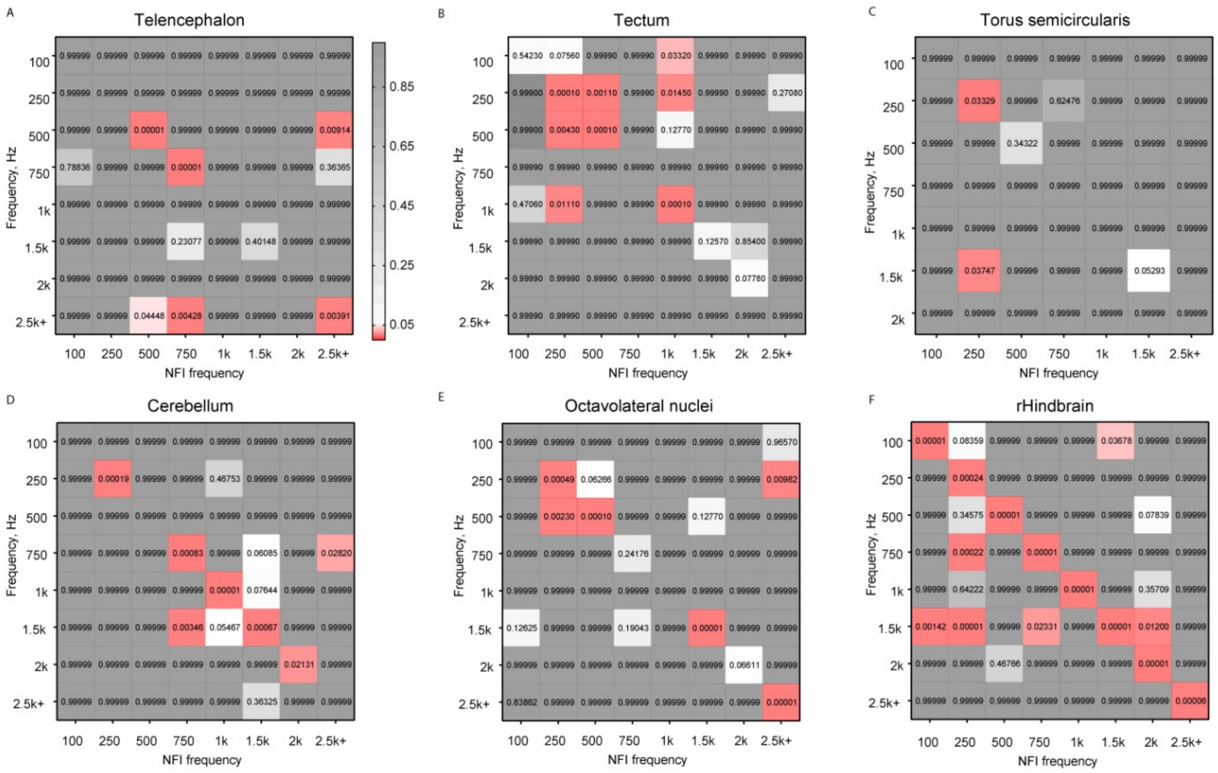


**Supplementary Figure 4 – P-values of the brain regions shown in figure 4** – P-values of the nearest neighbor analysis using the non-parametric Kruskal-Wallis test with Dunn's correction for multiple comparisons. Significant P-values ($P<0.05$) in red, and the remaining P-values as a gray gradient (bar in A).
